# E2EGraph: An End-to-end Graph Learning Model for Interpretable Prediction of Pathlogical Stages in Prostate Cancer

**DOI:** 10.1101/2023.03.09.531924

**Authors:** Wenkang Zhan, Chen Song, Supratim Das, Timothy R. Rebbeck, Xinghua Shi

**Affiliations:** Dept. of Computer & Information Sciences, Temple University, Philadelphia, PA, USA; Dept. of Biology, Indian Inst. of Science Education and Research - Pune, Pune, MH, India; Harvard TH Chan School of Public Health, Dana Farber Cancer Institute, Boston, MA, USA

**Keywords:** gene expression, cancer, disease prediction, graph attention network, interpretability

## Abstract

Prostate cancer is one of the deadliest cancers worldwide. An accurate prediction of pathological stages using the expressions and interactions of genes is effective for clinical assessment and treatment. However, identification of interactions using biological procedure is time consuming and prohibitively expensive. A graph is a powerful representation for the complex interactome of genes, their transcripts, and proteins. Recently, Graph Neural Networks (GNNs) have gained great attention in machine learning due to their capability to capture the graphical interactions among data entities. To leverage GNNs for predicting pathological stage stages, we developed an end-to-end graph representation and learning model, namely E2EGraph, which can automatically generate a graph representation using gene expression data and a multi-head graph attention network to learn the strength of interactions among genes and make the prediction. To ensure the reliability of model prediction, we identify critical components of graph representation and GNN model to interpret prediction results from multiple perspectives at gene and patient levels. We evaluated E2EGraph to predict pathological stages of prostate cancer using The Cancer Genome Atlas (TCGA) data. Our experimental results demonstrate that E2EGraph reaches the state-of-art prediction performance while being effective in identifying marker genes indicated by interpretability. Our results point to a direction where adaptive graph construction and attention based GNNs can be leveraged for various prediction tasks and interpretation of model prediction in a variety of data domains including disease prediction.

## I. Introduction

Cancer has been a common yet dangerous and lifethreatening disease of human beings. Extensive research has shown that early-stage cancer diagnosis predicts cancer treatment outcomes and improves survival rates. Therefore, early-stage screening and identifying cancer types have significant social and economic impacts. Massively parallel high-throughput sequencing technologies have generated a vast amount of genomics and transcriptomics cancer data. The development of a community resource project, The Cancer Genome Atlas (TCGA) [1], has generated comprehensive molecular profiles, including somatic mutation, copy number variation, gene expression, DNA methylation, microRNA expression, and protein expression of 33 different human tumor types, which provides with the ability to better predict cancer and understand the molecular basis of cancer etiology and treatment [2]. Therefore,

However, knowledge between molecular profiles and cancer is still limited, and identifying them by the biomedical procedure is time-consuming and prohibitively expensive. As a result, there is an increasing need to build efficient and feasible computation methods for cancer prediction especially leveraging the application of emerging machine learning techniques [3]–[8]. In the past few years, the prevalence of deep neural networks (DNNs) has led to more precise cancer diagnosis due to its ability of extracting high level features with meaningful connections among features [7]–[13]. Particularly, graph neural network (GNN) is a successful technique to iteratively learn complex relationships between objects by using the topology of graphs [14]–[19]. These GNNs are particularly powerful for learning graph representations toward better disease prediction by capturing the instinct relations from data. Recently, GNNs have achieved a superior success in a wide range of applications of biomedical data and deliver better results than competing deep learning methods. This makes GNNs an attractive tool for predicting cancer stages from gene expression data since the expression profiles of multiple genes in individuals are highly correlated, which can be reflected by complex relationships in the form of gene coexpression, regulatory networks, gene-gene interactions and protein-protein interactions. Hence, changes in the topology of the graph or just change in weights of the edges of the graph in a GNN can indicate changes in normal cellular mechanisms mediated by transcripts, genes, corresponding proteins, and their downstream biochemical pathways resulting cancer phenotype. It is thus straightforward to utilize graph representations to learn and represent such complex relationships in gene expression data in GNNs to improve prediction of disease risks and cancer stages.

However, one major challenge of implementing GNNs is to construct suitable graph representation that effectively embeds complex mechanisms and relationship of data. Existing methods in biomedical areas deploy statistical analysis, such as weighted gene co-expression network analysis (WGCNA) [20] and Pearson’s correlations [21]–[24], to measure the similarities among gene expressions and then transform the similarities into a graph representation to capture the relationships among genes. However, such an static strategy hardly reaches an optimal graph representation of genomics data and is usually conducted independently from predictive models. Therefore, remains a challenging problem to construct a powerful graph representation of data and includes this process into predictive modeling in an end-to-end optimization framework.

Apart from the prediction performance of cancer diagnosis and prognosis by computational techniques, it is an increasing concern to make the predictive model understandable and interpretable. Interpretability is a widely adopted method ensuring the reliability of a predictive model and the trustworthiness of its outcomes [25]–[27]. In biomedical and clinical applications, it is crucial to identify biological components such as genes and related biomarkers involved in the model outcome prediction and decision-making process. Additionally, a predictive model with outstanding prediction performances may have revealed patterns that might lead to new hypotheses and thus need to be interpreted and explained. Hence, it is non-trivial yet desirable to consider the interpretability problem of a predictive model to gain biomedical and clinical insights.

To handle the two aforementioned problems with an aim to enhance cancer early-diagnosis by capturing instinct relations among genes embedded by graphical computational techniques, we propose an end-to-end graph learning method (E2EGraph) for predicting cancer stages from patients’ genomics. E2EGraph integrates the construction of graph representation from genomics and knowledge learning to predict cancer stages empowered by GNNs.. In detail, we first compute the co-effect of genes with pairwise dot product. Then, we employ a differential threshold adapter with a parameterized threshold to transform the values of dot product into edges of a graph. This threshold can be optimized in each training epoch with classifier. With the graph representation constructed this way, a multi-head graph attention network (multi-head GAT) is utilized for predicting pathological stages as a classification task. Additionally, to ensure reliability of E2EGraph, we identify critical components of graph representation and enable multi-head GAT to interpret prediction results from multiple perspectives, including at gene, gene-gene interaction, and individual patient levels. The innovative contributions of this paper can be summarized as follows:

### Adaptive graph learning

To find an appropriate and efficient graph representation of genomics, we proposed a novel graph construction method that captures the combinatorial effects of multiple genes in graph. In detail, after computing the pairwise dot product co-effect value derived from the expressions of any pair of genes, we dynamically construct graph representation of gene expression quantification using an adaptive threshold which can be optimized during training process. This dynamically updated threshold for setting edges in each training epoch enable finding an optimal graph to facilitate downstream cancer prediction in E2EGraph.

### Multi-head attention mechanism

To improve the overall representation capability and prediction performance of E2EGraph, we employed a multi-head attention mechanism into a GNN model. The multi-head attention module repeats computation of multiple attentions independently and then combines all attention scores for final prediction. By doing this, the GNN model in E2EGraph is focused on the important portions of the graph network and outputs a better prediction of corresponding cancer stages of patients.

### Interpretability

An important perspective of building powerful predictive models for precision medicine is that the prediction is explainable and results are interpretable. Hence, we implement state-of-the-art intepretable machine learning techniques to provide the interpretility of our models as well as the data. Specifically, to understand E2EGraph’s mechanism, we employed gradient saliency and attentions to explain the knowledge (gene importance and their co-effects) that E2EGraph learned from a given dataset. In the data persepective, we leverage local interpretable model-agnostic explanations (LIME) [26] to analyse the effects of noise in gene expression to cancer stage prediction.

### End-to-end learning

To the best of our knowledge, E2EGraph is the first work that provides an end-to-end learning framework seamlessly combining graph construction and graph neural network learning for omics data analysis. Unlike alternative methods where a fixed graph threshold was used the adaptive threshold learning enables the updating of graph representation when fitting the model so that it can unified to optimize graph representation as well as graph network. Our experimental results in prostate cancer stage prediction from gene expression data, showed that E2EGraph is effective and has the potentials to be generalized to other types of data.

## II. Method

The code and results of this study is publicly available at Github [28]. In this section, we describe the proposed end-to-end graph learning model (E2EGraph) that consists of one graph representation construction module, one knowledge learning module, one prediction module and one interpretable module (**Fig. 1**). More specifically, given gene expression profiles of patients, we first measure the co-effect of two genes with dot product [29] of corresponding gene expression value. We compute all of pairwise co-effect values and set edges between those nodes whose co-effect values are greater than a threshold. After representing graph, we employ a multi-head graph attention network (multi-head GAT) to learn hierarchical node features. At the output sides of multi-head GAT, we concatenate the input graph and the hierarchical graph to a prediction head for outputting prediction probabilities. Meanwhile, we also provide explanations at gene, sub-network and patient levels to interpret our model predicting using gradient saliency, attention visualization, and LIME methods accordingly.

**Fig. 1.**
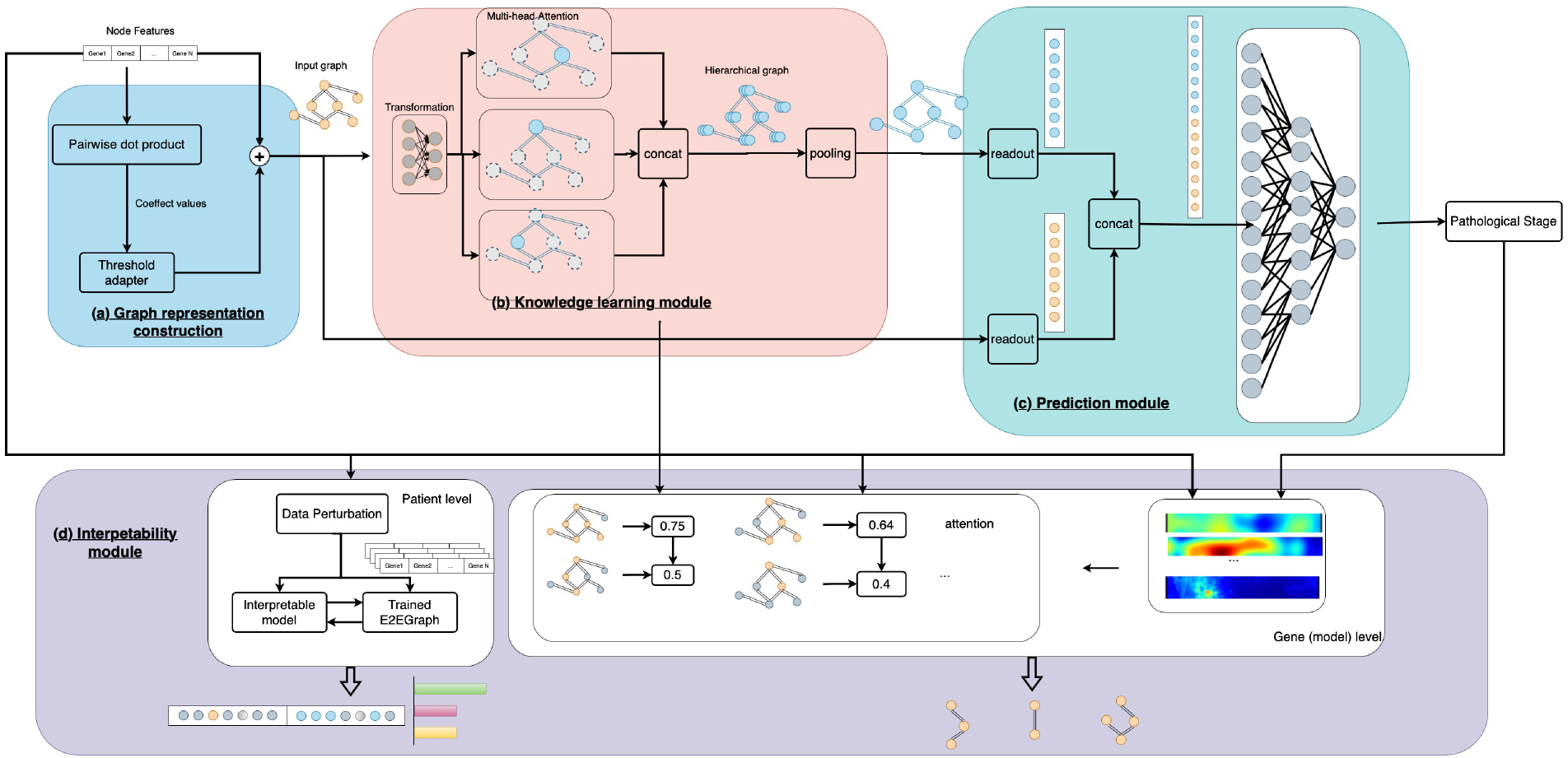
Overview of the end-to-end graph learning model (E2EGraph model): (a) Graph construction module. Firstly, the pairwise co-effect of two genes are computed. Then edges between two genes are constructed if the co-effect value being greater than the threshold value. A threshold adapter is employed to update the threshold value during training. (b) Knowledge learning module. We deployed a multi-head attention graph network for capturing graph knowledge. Multiple independent attention heads compute parallelly in this module and then generate the hierarchy graph by concatenating outputs from each head. (c) Prediction module. We utilized the original graph and the hierarchy graph for final prediction. After concatenating two graphs, a prediction head with three linear layer is deployed to generate prediction probabilities. (d) Interpretability module. We averaged the gradient saliency and attentions of all patients to explain genes expression and their co-effects learned by E2EGraph from the whole dataset. Besides, we employed LIME to explain the influence of noise in gene expression to the cancer stage of an individual patient.

### A. Graph construction

In this paper, we treat each gene position as a node and construct graph based on the relationship among nodes. Assuming that there should be an edge if two nodes effect together strongly, we connect edges depending on a quantitative co-effect value. In detail, we first compute the pairwise co-effect value and then we set edges between those pairs of nodes that have a co-effect value greater than a threshold.

In mathematical representation, given a *K* dimensional numerical genomics denoted as *V*, our goal is to construct a graph representation which is specified by nodes V and edges **E**. We compute dot products *s_ij_* = *v_i_v_j_* among any pair of nodes to measure the combinatorial expression, where *i, j* ∈ {1, 2,…, *K*} represents any two nodes and then obtain a matrix of dot product **S** ∈ *R^K×K^*. By setting a threshold t, the edges **E** can be found at **S** ∈ *R^K×K^* ≥ *t*

It is noticed that if the t is set manually and to be fixed in the graph knowledge learning, it is hard to say the graph representation is suitable. Although in the alternative state-of-art employed multiple strategies, for instance, cross-validation, to find a suitable threshold, it is still difficult to obtain a optimal solution. In this regard, we novelly designed a threshold learning algorithm and try to use this algorithm to learn the optimal threshold value t. In detail, we introduced a differentiable adapter with a parameterized threshold to transform the values of dot product into edges **E***^K×K^* of a graph as follows.

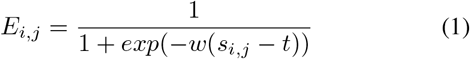

where *t* denotes the parameterized threshold, *w* represents a scalar forcing *E_i,j_* to neighbourhood of 0 or 1, and *E_i,j_* represents the edge between *v_i_* and *v_j_*.

The threshold adapter is embedded to the graph network and is to be trained with graph network. In this way, it enables the model to generate the optimal graph representation in each training epoch.

### B. Knowledge learning

After constructing graph representations **G** = {*G*_1_, *G*_2_,…, *G_n_*} for patients, where *N* denotes the total number of patients in the dataset, our goal is to learn the knowledge not only from the features themselves but also from the topological information of the graphs. To do so, we employ a graph attention network (GAT) to learn hierarchy node features from a weighted co-effect of genes realized by attention mechanism.

To begin with, we perform feature transformation *F*(.) to every node (gene) using a same linear function followed by a nonlinear activation *F* (*v_i_*) = *σ*(*Wv_i_*) where *W* represents weights of the linear function, and *σ*(.) represents the nonlinear activation. After feature transformation, we update the node features by aggregating messages from neighbouring nodes of *v_i_* by attention mechanism.

To introduce attention, we first compute coefficients of a node to its neighbours *e_i,j_* = *a*(*Wv_i_*∥*Wv_j_*) where *a* is a feedforward neural network, ∥ represents concatenation. and *e_i,j_* indicates the importance of *v_j_* to *v_i_*. Then we normalize the attentions

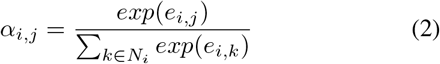

With a purpose to stabilize the learning process and introduce attention power, we independently compute multiple attention heads in a parallel manner and then concatenate their outputs to generate a hierarchy graph.In this way, the node features of the hierarchy graph is updated as

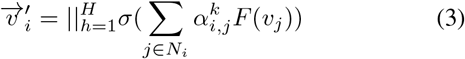

where 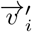 is hierarchy features of node *i, H* is the number of attention heads, ∥is concatenation, 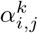 are the attention coefficients of the *k*-th attention mechanism.

### C. Prediction

The knowledge learning module produced a hierarchy graph of gene expression. In the prediction module, we use the original graph and the hierarchy graph for final prediction. To make these two graphs fit into the prediction head, we first reduce the hierarchy features to one dimension by pooling operation. Then, we vectorize the nodes of these two graphs by concatenation as a vector *z* ∈ *R*^2*K*^. Then, we employ prediction head with three linear layers to generate predition probabilities.

In the training process, we measure both the graph representation construction and the prediction by a loss function

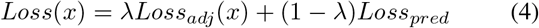

where *λ* is in the range of (0, 1). To be specific, prediction result is measured by cross-entropy and graph representation construction is measured by 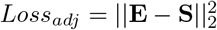.

### D. Interpretability

We employ multiple state-of-the-art methods for interpretable machine learning to provide interpretability of the prediction mechanism of E2EGraph and the effect a gene to its corresponding prediction.

#### 1) Extraction of important genes for model prediction

To interpret the prediction mechanism of E2EGraph, we employ gradient saliency to visualize the weights that E2EGraph assigns to each gene for prediction. Assuming that we have an prediction *y_i_* of the *i*-th patient, the graident saliency of y_i_ to node (gene) *p* is 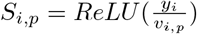. Then, we average gradients of every node (gene) with respect to all of patients by 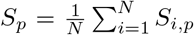 Picking those highest gradient scores, we can find the nodes (genes) which are most important to working mechanism of E2EGraph. Besides, we visualize the attention scores of E2EGraph to the genes to explain the co-effect to the E2EGraph mechanism.

#### 2) Interpretation of cancer stage prediction for an individual patient

To facilitate precision medicine, it is desirable that can explain the prediction of a patient’s cancer stage based on his/her gene expression profiles.

To do so, we employ LIME method to explain the prediction of an individual patient [26]. Given the gene expression *x* of a patient, LIME generates a sub dataset *D* by perturbing x. With *D,* LIME then trains an interpretable model *g* by comparing the output *y* of the trained E2EGraph model *f* based on *D*. We find an optimal interpretation *g\** by minimizing the loss

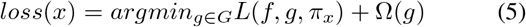

where Ω(*g*) is model complexity, *G* is the family of possible explanations, and *π_x_* defines the neighborhood around *x*.

## III. Experimental setup

We demonstrated the application of E2EGraph to provide interpretable prediction of prostate cancer stages using gene expression profiles in the TCGA data. We compared the performance of E2EGraph with other methods including logistic regression (LR), multi-layer perceptron (MLP), convolutional neural network (CNN), multi-Level Attention Graph Neural Network (MLA-GNN) (the state-of-the-art model on genomics using handcraft threshold for graph construction), using evaluation metrics of accuracy, precision, recall, and F1 score.

### A. Datasets

We collected High-Throughput Sequencing - Fragments Per Kilobase of transcript per Million mapped reads (HTSeq-FPKM) as gene expression quantifications and clinical information of each patient with prostate gland cancer from The Cancer Genome Atlas (TCGA) [1] using the GDC Data Transfer Tool [30]. These HTSeq-FPKM values were then normalized to a range of [0, 1] with min-max normalization. From the clinical information, we extracted pathological stages of patients according to the American Joint Committee on Cancer (AJCC) TNM staging system. To ensure that we have the most available data without worries about population stratefications, we used only patients with European ancestry. To mitigate severe unbalanced number of samples in different pathological stages of prostate cancer, we neglected pathological stages I and only use pathological stages II, III, IV as labels, which contains 130, 62 and 47 patients respectively. Besides, to reduce the high dimensionality of gene expression, we selected most frequently mutated genes provided by TCGA data portal [1]. In total, our dataset include the quantifications of 538 genes in 244 patients. To validate the effectiveness and robustness of the proposed method, we conduct 10-fold cross-validation in the experiments.

### B. Implementation details

For logistic regression (LR) and multi-layer perceptron (MLP), we don’t perform any further processing to both of the training set and testing set. The number of hidden units in hidden layers of MLP is set to {512, 256, 64, 3}.

For convolutional neural network (CNN), we transform the gene expression quantification of each patient into a 24 × 24 two dimensional (2D) array, by filling 0s to fit into the 2D format [31]. We employ a 2D convolutional layer with output channel of 8, kernel size of (3, 3), stride of (3, 3), a ReLU activation, and a max pooling layer with kernel size of (2, 2), stride of (2, 2), followed by fully connected layers with units of {64, 32, 3}.

For MLA-GNN, the state-of-the-art GNN methold applied on genomics, we follows the setting of [32] by constructing graph representation with weighted correlation network analysis (WGCNA) based on the training set.We select the threshold of WGCNA similarity for constructing the adjacency matrix with grid search strategy from {0.5, 0.6, 0.7, 0.8, 0.9}.

For E2EGraph, we employ one GAT layer with hidden units of 8, and predict head with units of {512, 128, 3}. We implement all of the experiments with Pytorch. For all of the above methods, we set learning rate to 0.001, training epoch to 200, batch size to 32 and employ the Adam optimizer to optimize the parameters.

## IV. Results

### A. Prediction performance

The performance of prediction of pathological stages in prostate cancer is evaluated by metrics including accuracy, precision, recall, F1 score. Table III-B showed the comparison of the performance among LR, MLP, CNN, MLA-GNN, and E2EGraph. In general, GNN models (MLA-GNN, E2EGraph) trained on graph representations of gene expression have better performance than other machine/deep learning models (LR, MLP and CNN), suggesting that underlying combinatorial effects such as epistasis, are attributed to the prediction of pathological stages in prostate cancer. Nevertheless, CNN does not see improvements over LR and MLP, indicating that combinatorial effects among gene expression are non-Euclidean based and thus extracting information from nearby genes is less meaningful. Hence, we showed that constructing graph representation using statistical analysis is beneficial to capture the combinatorial effects of gene expression and help with predicting prostate cancer stages.

Our proposed E2EGraph method reaches an outstanding performance when evaluated by accuracy (0.7000), precision (0.7000), and F1 score (0.7000), and outperformed MLA-GNN at recall (0.6887 vs. 0.6546). These results suggested that the adaptive threshold for graph construction of E2EGraph have the capacity of constructing better graph representations for prediction through the end-to-end training procedure.

### B. Interpretation of prediction results

Besides its outstanding performance, E2EGraph provides interpretations of prediction results at gene, sub-network and patient levels using different interpretable strategies.

#### Interpretation of E2EGraph for predicting pathological stages

We utilized the gradient saliency of every individual patient in testing set to find the most influential genes for predicting stages in prostate cancer using E2EGraph. Fig. 2 shows the top 30 genes in the testing dataset that are critical for predicting prostate cancer stages using E2EGraph, meaning that E2EGraph focus on these 30 genes most when making prediction. Among the 30 genes we found to be most important in making predictions of prostate cancer stages in E2EGraph, some of the most interesting ones are BMP5, GRM3, CCND2, RSPO2, PDE4DIP, CDK12, WT1, ETV4, PMS1, and ZNRF3. Abnormal expression caused by increased copy numbers of BMP5 has been reported in prostate cancer compared to benign prostatics issue [33]. SNP and gene-based association analysis of the RTK/ERK pathway has found that reduced expression of CCND2 promoted cell proliferation and its overexpression inhibited cell growth of prostate cancer [34]. ETV4 fusions with KLK2 and CANT1 are reported as predominant common characteristics androgen-induction and prostate-specific expression [35]. Genomic alterations in CDK12 are associated with metastatic castrate-resistant prostate cancer [36]–[38]. GRM3 is observed to have strong accumulation effects on overall survival in renal cell carcinoma [39].

**Fig. 2.**
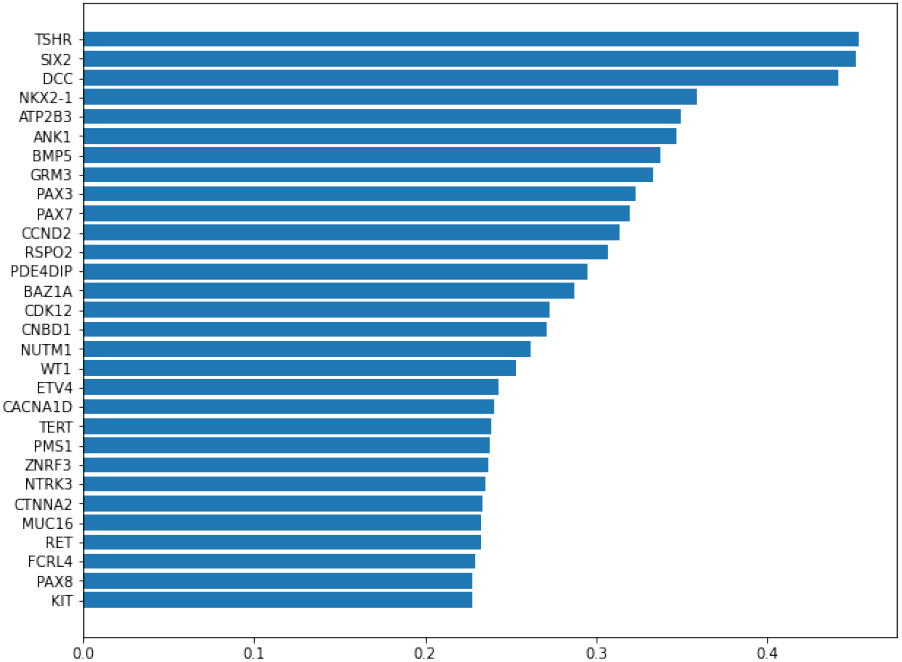
Visualization of the Top 30 important genes (Y axis) and their importance on the test dataset ranking by the gradient saliency values of these genes (X axis).

Using ShinyGO [40], we performed Gene Ontology (GO) enrichment analysis of the 30 genes for predicting prostate cancer stages. GO analysis results suggest that most of these genes are related to hemi-methylated DNA-binding, P-type calcim transporter activities and protein-hormone receptor. (**Fig. 3**).

**Fig. 3.**
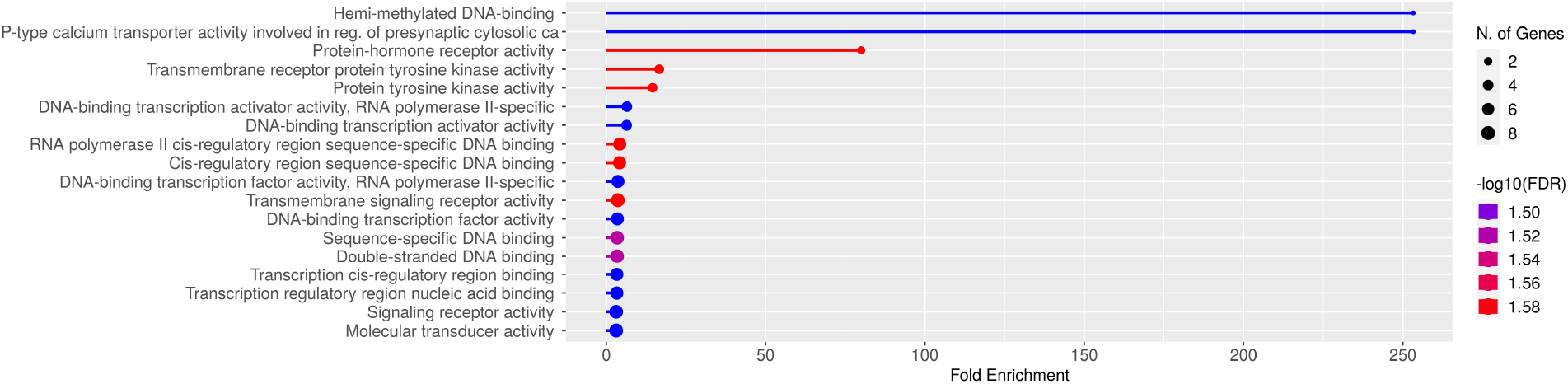
Gene Ontology (GO) enrichment of top 30 important genes. X-axis represents the fold enrichment of certain GO terms with respect to our list of genes, while the color gradient depicts the false discovery rate (FDR) of corresponding GO terms. Most of these genes are related to DNA damage repair, DNA damage checkpoint signaling pathways, and developmental pathways.

**TABLE I.**
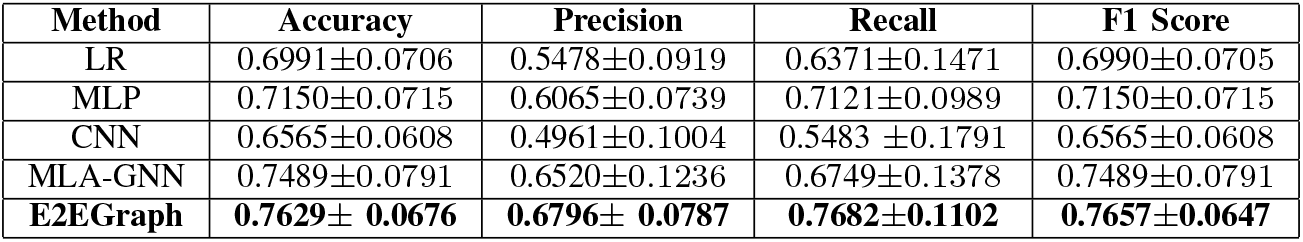
Performance of multiple models in comparison with that of our E2EGraph model for prediction of prostate cancer stages

Besides intepreting individual genes, we analyze the combinatorial effects among genes for their prediction of prostate cancer stages by visualizing the attentions on the top 30 genes in E2EGraph (**Fig. 4 (a)**) and visualizing their protein products and the protein-protein interactions (PPI) using STRING database [41] (**Fig. 4 (b)**). The attention mechanism of E2EGraph successfully learn part of the interactionschen: in which espect?.

**Fig. 4.**
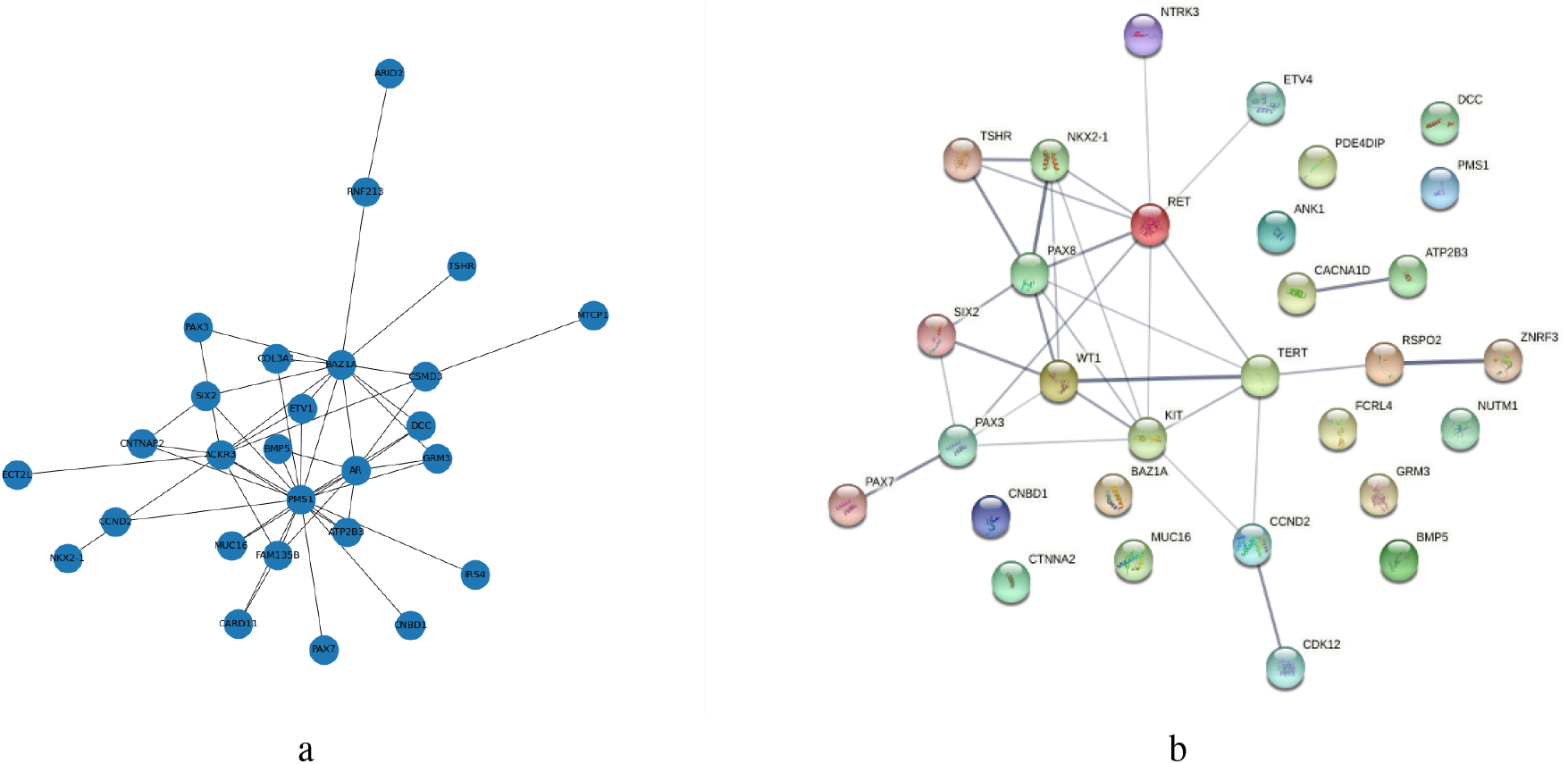
Interactome sub-graph visualization. (a)**Visualization of the attention mechanism among top 30 genes derived by E2EGraph:** Each Node represents a gene. While the width of the edges represents the strength of relation among those genes. (b)**Visualization of protein protein association graph of gene product of the same 30 genes from E from STRING Database:** Here each node represents protein of that gene. Cyan and red lines represents known interaction from curated database and experimentally determined interactions. The Black line represents co expression of those gene products.

Our study also finds a less obvious gene, like BMP5, GRM3 also is not connected in the protein-protein interaction graph of the STRING database (**Fig. 4 (b)**), but in the graph generated by E2EGraph, this gene seems to be connected with many other genes. As this gene has not yet been experimentally associated with prostate cancer, this result might be of interest for biochemical validation.

#### Interpretation of the prediction of a patient’s cancer stage

We employ LIME to give a probability for a patient’s cancer stage based on his gene expression using E2EGraph, and identified important genes by tweaking the feature and see its impact on prediction. We randomly chose to analyze the prediction of the 3-rd patient of the dataset with its prediction as pathological stage IV (**Fig. 5**). Another relevant gene product was SMARcD1; this protein is a subunit of the SWI/SNF complex, which affects gene expression regulation by ATP-dependent chromatin remodelling [42]. Researchers have connected upregulation of SMARcD1 with de-differentiation of committed basal cells of prostate gland to stem cells [43].

**Fig. 5.**
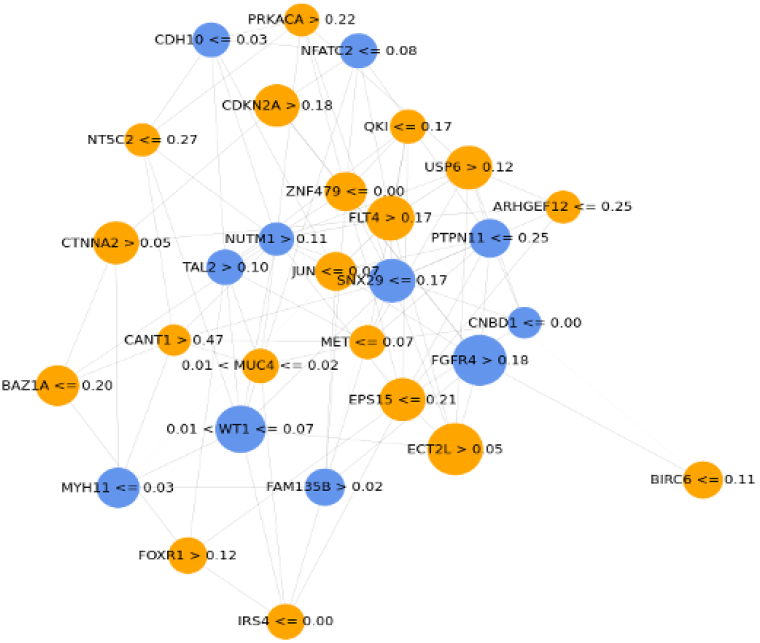
Visualization of the Top 30 important genes and their relations by LIME and attention.

Although most of these genes are related to their function inside the nucleus, SDC4 is a transmembrane proteoglycan that mediates the interaction of cell with its outer environment. As SDC4 plays an important role in *α*5*β* 1 integrin-dependent focal adhesions, organization, and cell migration, which makes it an interesting gene for studying malignancy of cancer tissue [44]. Although this gene has not yet been connected with prostate cancer, the literature suggests downregulation of SDc4 in malignant breast tissue with respect to normal breast cancer tissue [44].??

Genes found to drive the prediction of an individual patient are equally relevant. In every instance of a randomly selected patient, we find different sets of genes having more effect in making the prediction. This suggests that various prostate cancer phenotypes might rise from perturbation of completely different pathways. For patient 3, we find one such interesting gene LASP1 whose expression quantification drives the prediction of this patient’s cancer stage. LASP1 has already been suggested to be a promising drug target, as knockdown of LASP1 in prostate cancer cell lines has shown to decrease cell proliferation and migration significantly in an NF-κB dependent manner [45]. The same patient also has MACC1 as an important driver of prediction. Interestingly this gene has also been shown to inhibit NF-κB nuclear translocation [?]. Another such gene for this patient is EED, which also interacts with NF-κB transcription factor [47]. We see multiple top genes driving predictions connected with the NF-κB pathway, which plays a key role in any cancer as this affects pathways controlling apoptosis and cell migration [48], a treatment targeting NF-κB can be prioritized. Researchers have already identified NF-κB as a very potent drug target [48].

These results suggest that some of the genes identified by our in-silico methods have biological relevance, which is established by experiments. Thus genes found in our method might be of interest for further experimental validation, which can be potential drug targets or prognostic markers. Our case study on one patient also suggests the applicability of our model in precision medicine, as the top 10 driver genes might allow physicians to prioritize drug targets and personalize treatment strategies for this patient based on his gene expression profile.

## V. Conclusion

In this paper, we propose an end-to-end graph learning (E2EGraph) model for prediction of pathological stages in prostate cancer. To capture the relationships, we construct graph representations by using pairwise dot product to measure the combinatorial effects among genes and developing an adaptive threshold to transform the values of dot product into adjacency matrix. With the constructed graph, we employ graph attention network for knowledge learning from the graph representation. Finally, we employ an MLP classifier to predict the pathological stage in prostate cancer. In addition to the prediction, to make the prediction faithful, we deploy gradient saliency, attention visualization, and local interpretable model-agnostic explanations (LIME) methods to explain the graph representation as well as the E2EGraph model at gene, subnetwork and patient levels. Experimental results show that our E2EGraph model is effective in interpretable prediction of the pathological stage in prostate cancer with TCGA dataset and has the ability to identify marker genes and discover combinatorial effects. The E2EGraph has the potential to be adapted for predicting other types or disease such as lung caner, as well as using other types of data, e.g. DNA methylation, protein quantifications.

We also checked the biological relevance of 30 genes found most useful in prediction of prostate cancer stages. Many of those genes have already been associated with prostate cancer or other cancers, while some are not yet associated with cancer pathways. Although, in this study, we apply our model to transcriptomics data of prostate cancer, our results suggest the applicability of our model in interpretable data mining for finding drug targets and prognostic markers for other types of datasets like genomics, epigenomics and proteomics data and for other complex diseases as well. We have also demonstrated applicability of our model in precision medicine with a case study on predicting the pathological stage of one patient based on the patient’s gene expression.

Although the proposed E2EGraph model finds an optimal solution for graph representation construction and achieves outstanding performance. However, there are some aspects needed to be improved in future work. Firstly, although graph neural networks have shown potential in omics datasets by outperforming other machine learning methods and deep neural network methods, a better design of graph neural network algorithms is required and can further improves model performance. Besides using molecular data in this study, other modality of data such as medical images should be included into a multi-modal method to improve the prediction tasks of diseases like cancer. Secondly, it is still prohibitively expensive or sometimes impossible to collect biomedical data that is not biased and covers all population or disease subtypes. In this regard, we will focus on knowledge transferring by transfer learning techniques in future disease prediction.

